# Intraspinal stimulation with silicon-based 3D microelectrode array for bladder voiding in cats before and after spinal cord injury

**DOI:** 10.1101/2020.01.13.905364

**Authors:** Victor Pikov, Douglas B. McCreery, Martin Han

## Abstract

Bladder dysfunction is a significant and largely unaddressed problem for people living with spinal cord injury. Intermittent catheterization does not provide volitional control of micturition and has numerous side effects. Targeted electrical microstimulation of the spinal cord has been previously explored for restoring such volitional control in the animal model of experimental spinal cord injury. In this study, the development of the intraspinal microstimulation array technology was continued and evaluated in the feline animal model for its ability to provide more focused and reliable bladder control after a complete spinal cord transection. For the first time, the intraspinal multisite silicon array wad built using novel microfabrication processes to provide custom-designed tip geometry and 3D electrode distribution, better cost efficiency, reproducibility, scalability, and on-the-probe integration of active electronics. Long-term implantation was performed in 8 animals for a period up to 6 months, targeting the dorsal gray commissure area in the S1 sacral cord that is known to be involved in the coordination between the bladder detrusor and the external urethral sphincter. About one third of the electrode sites in the that area produced micturition-related responses. The effectiveness of stimulation increased starting from one month after spinal cord transection (as evaluated in one animal), likely due to supraspinal disinhibition of the spinal circuitry and/or hypertrophy and hyperexcitability of the spinal bladder afferents. Further studies are required to assess longer-term reliability of the developed intraspinal microstimulation array technology in preparation for eventual human translation.

## Introduction

Approximately 300,000 individuals are currently living with spinal cord injury (SCI) in the United States with annual incidence of 54 cases per one million, or about 17,700 new cases annually (NSCISC 2018). Restoration of bladder and bowel dysfunctions has the greatest priority for people living with SCI, higher than regaining walking ability (Anderson 2004). Intermittent catheterization is presently used for emptying the paralyzed bladder, but that procedure has numerous side effects, most notably, the development of urinary tract infections and chronic cystitis (Anderson et al. 2019, Roth et al. 2019, Tofte et al. 2017). Furthermore, intermittent catheterization does not address the underlying loss of volitional control of micturition. The approach based on electrical microstimulation inside the spinal cord may provide such volitional control. In our earlier study using the cat model of SCI, targeted electrical microstimulation in the dorsal gray commissure (DGC) was shown to be effective in triggering physiologically coordinated bladder contraction and relaxation of the external urethral sphincter (EUS) (Pikov, Bullara, and McCreery 2007). The microstimulation-induced voiding was elicited in the majority of the animals (15 out of 22) for a period up to 14 months although the voiding efficiency was rather low in most animals. Twenty animals in that study were implanted with a microwire array with two-dimensional electrode distribution, while only two had the multisite silicon array with 3D electrode distribution. We hypothesized that 3D electrode distribution would provide better access to the spinal gray matter nuclei, particularly in the DGC area and, therefore, would facilitate better targeting with lower current levels and reduced current spread to unrelated nearby neuronal groups (Krucoff et al. 2016, Giszter 2015). Other studies have attempted to achieve neuroprosthetic bladder control with individual spinal microwires (Tai et al. 2004), spinal microwire arrays (Gaunt et al. 2006), spinal epidural stimulation (Gad et al. 2014, Harkema et al. 2011, Abud et al. 2015, Pettigrew et al. 2017, Woellner, Krebs, and Pannek 2016), and stimulation of the spinal, pudendal, and pelvic nerves (Chew et al. 2013, Peh et al. 2018, Yoo et al. 2007, Tai et al. 2011). Intraspinal microstimulation following SCI has also been used for restoring limb function, including microwires (Giszter et al. 2000, Mushahwar et al. 2002, Mushahwar and Horch 2000, Dalrymple et al. 2018, Moritz et al. 2007, Sunshine et al. 2013, Mercier et al. 2017, Bamford and Mushahwar 2011), microwire arrays (Holinski et al. 2016, Saigal, Renzi, and Mushahwar 2004, Grahn et al. 2016, Kasten et al. 2013), and multisite silicon arrays (Zimmermann, Seki, and Jackson 2011, Borrell et al. 2017). Comprehensive reviews have been published describing various neuroprosthetic approaches for restoring the bladder control after SCI (McGee, Amundsen, and Grill 2015, Bamford and Mushahwar 2011, Pikov 2008, Gaunt and Prochazka 2006) and for restoring limb function after SCI (Giszter 2015, Ievins and Moritz 2017, Gaunt et al. 2006, Mushahwar et al. 2007).

The goal in this study was to develop the next generation of chronically-reliable multisite stimulating probes, to optimize the methods of assembling these probes into 3D arrays, and evaluate the functionality of embedded electronics in the array. For the probe fabrication, we chose to use the deep reactive ion etching (DRIE) process, which allows a probe thickness of 100 μm (or more) and custom-designed tip geometry for more reliable insertion through thick pial membrane that covers the dorsal surface of the spinal cord. This is in contrast with a probe thickness of 15 μm fabricated using the wet etching process for our previous study (Pikov, Bullara, and McCreery 2007), which likely was too fragile for chronic implantation in primates without using a cumbersome mounting device for insertion (Sakamoto et al. 2009) or increasing the probe thickness to 80 μm without custom-shaping the tips (Ruther et al. 2010). The 3D microfabrication technology is potentially superior to 2D microwire fabrication by facilitating computer-aided design of the probe and tip geometry, better cost efficiency, reproducibility, scalability, on-the-probe integration of electronics, and reducing the tissue microtrauma caused by an array insertion due to a smaller number of shanks in multi-site probes as compared to single-site microwires with a similar density of electrodes in the DGC area (Pikov, Bullara, and McCreery 2007). Since the proposed 3D array contains 64 electrode sites on 2 probes, its connection via a 64-wire subcutaneous cable could be rather bulky and inflexible. In order to reduce its bulk and thus minimize the transmission of traction and torsion from the cable to the array and reduce the risk of tissue damage during long-term implantation, we employed the demultiplexing ASIC chip embedded on the probe. Optimization of the array design was done by assessing possible failure modes in the array prototypes during their semi-chronic implantation (up to 6 months). At the end of the study, we achieved an improved array design, and further studies would be required to assess its longer-term reliability during one-year (or longer) implantation time.

## Methods

### Probe fabrication

The probe consisted of eight 50-μm wide shanks spaced 300 μm apart (center-to-center), with each shank containing four 20×100 μm stimulating sites also spaced 300 μm apart (center-to-center) for equal site distribution in the transverse spinal cord plane dimensions. Layout of the bonding pads in the probe superstructure allowed the probes to be either wedge-bonded to the subcutaneous cable directly or wedge-to-ball-bonded through the demultiplexing ASIC chip (Figure 1).

**Figure 1.**
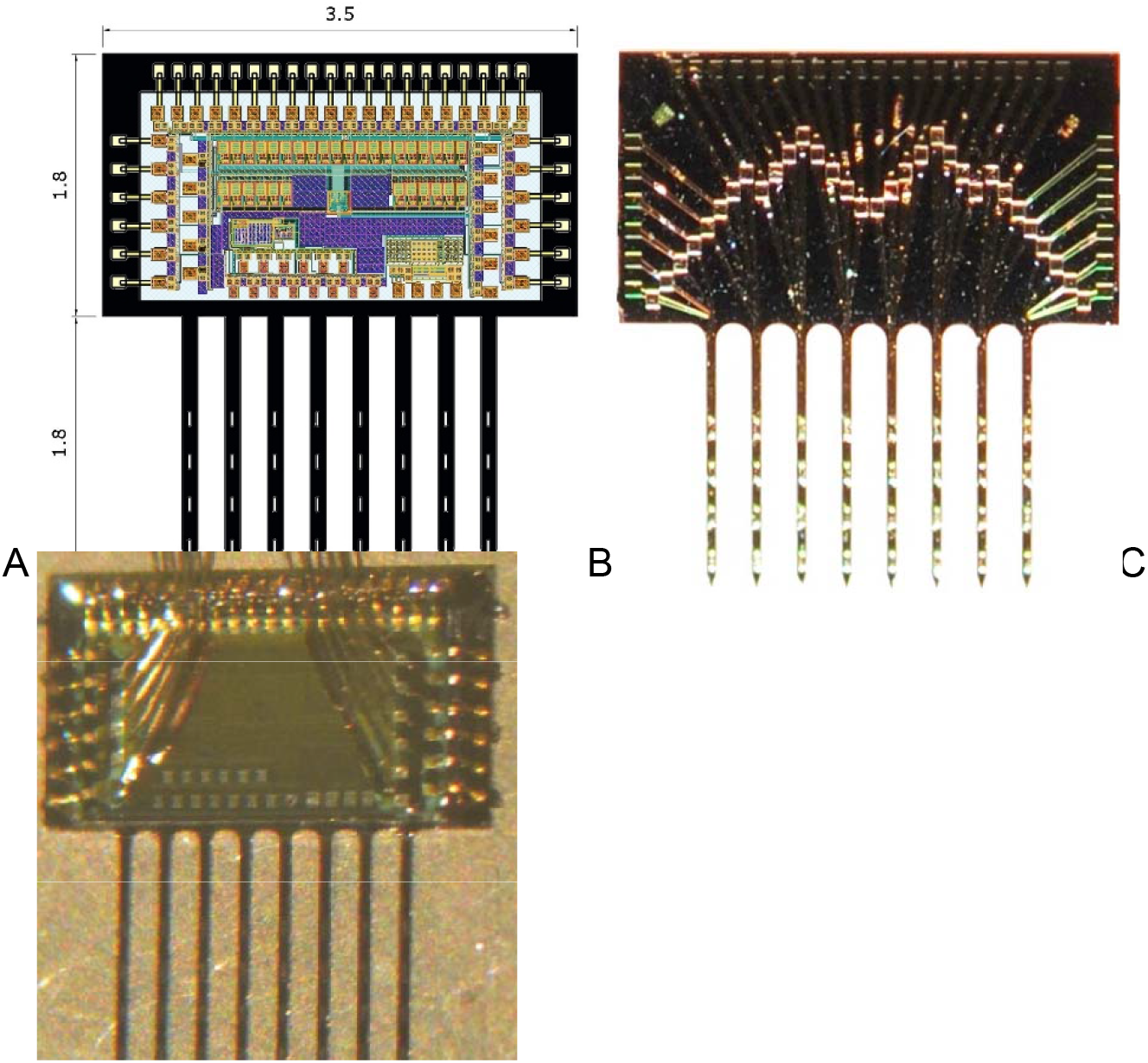
A) The probe layout with a wire-bonded ASIC chip (dimensions in mm); B) photo of a probe without the ASIC chip; and C) photo of a probe with the ASIC chip connected to 10 wires of the subcutaneous cable.

The probes were fabricated at the Micromachining laboratory at the California Institute of Technology according to the steps previously described (Han et al. 2011). Four-inch diameter silicon-on-insulator wafers were used, and the main insulation layers were silicon dioxide-silicon nitride-silicon dioxide. The tips of the probes were mechanically ground in order to reduce tissue dimpling and facilitate insertion into the spinal cord (Han et al. 2011). IrOx (iridium oxide) was electroplated onto the gold electrode sites to form electrodeposited IrOx film (EIROF), which increased the sites’ charge injection capacity and resistance to corrosion and dissolution during stimulation, compared to pure iridium and platinum (Robblee, Lefko, and Brummer 1983). The EIROF method was adopted from the EIC Laboratories (Norwood, MA) (Meyer et al. 2001). Briefly, the EIROF solution contained 0.5% oxalic acid and 0.13% iridium tetrachloride in 0.1M phosphate-buffered saline (PBS), and electrodeposition used the triangular potential cycling at 50 Hz and a sweep rate of 50 mV/s between potential limits of 0.6V to 0.8V versus Ag/AgCl using the potentiostat (PC4/300, Gamry Instruments, Warminster, PA) and the cyclic voltammetry software (CV package of Gamry Framework, Gamry Instruments) (Han, Liu, Bullara, Lossinsky, et al. 2004, Han, Liu, Bullara, Pikov, et al. 2004, Han et al. 2005). The stimulation sites had the dimensions of 20 × 100 μm (width × length) for a geometrical area of 2,000 μm^2^. Controlled current pulsing for determining charge injection density was done using charge-balanced pulses at an amplitude of ± 50 μA. This produced a total potential excursion of <= ± 0.9 V versus Ag/AgCl. The stimulus was administered using a custom stimulator and a control program written in QuickBasic.

### ASIC chip fabrication and assembly on the probe

The demultiplexer ASIC chip was custom-designed to our specified dimensions of 2.8 mm × 1.5 mm by Prof. Wentai Liu at the University of California Los Angeles. Using this 32-to-5 demultiplexer, the stimulation waveform can be routed sequentially to each of 32 sites in the interleaved mode using 5 digital address lines controlling a bank of analog switches. In addition to routing the stimulation waveform in vivo, the chip was also used in vitro during EIROF deposition process. The ASIC chip was first glued onto the probe with epoxy (EpoTek 301, Epoxy Technology, Billerica, MA). The wire bonds between the bonding pads on the demultiplexer and 3 edges of the probe were made with 25 μm-thick stress-relieved gold wire using wedge-to-ball bonding (Model 747677E, West Bond Inc, Anaheim, CA), as shown in Figure 3C. In addition, ten 90% platinum / 10% iridium wires with gold-plated ends from the subcutaneous cable were attached to 10 pads on the chip (Figure 1C) to provide 5 address lines, two supply voltages (positive and negative), ground, output bias, and input signal from the stimulator, which were bonded to pads on one edge of the chip and directly into a subcutaneous cable. The output cathodic bias of +400 mV was routed inside the chip to each of the 32 stimulating electrode sites in order to maximize the charge capacity of EIROF-deposited IrOx film on the sites during pulsing in vivo (Meyer et al. 2001). Connection between the probes and the head-mounted connector was via a subcutaneous cable (70 cm). The cable from the external controller contained 33 perfluoroalkoxy-isolated wires (outer diameter (OD) = 0.063 mm, 42 AWG, Cooner Wire, Chatsworth, CA), which were helically wound using a custom-made winding machine. The 20-wire cable was used for connecting to both probes (10 wires per probe) via the demultiplexer ASIC chip, while the 33-wire cable was used for connecting to a single probe without the chip (one of the wires was connected to a platinum ground plate).

**Figure 3.**
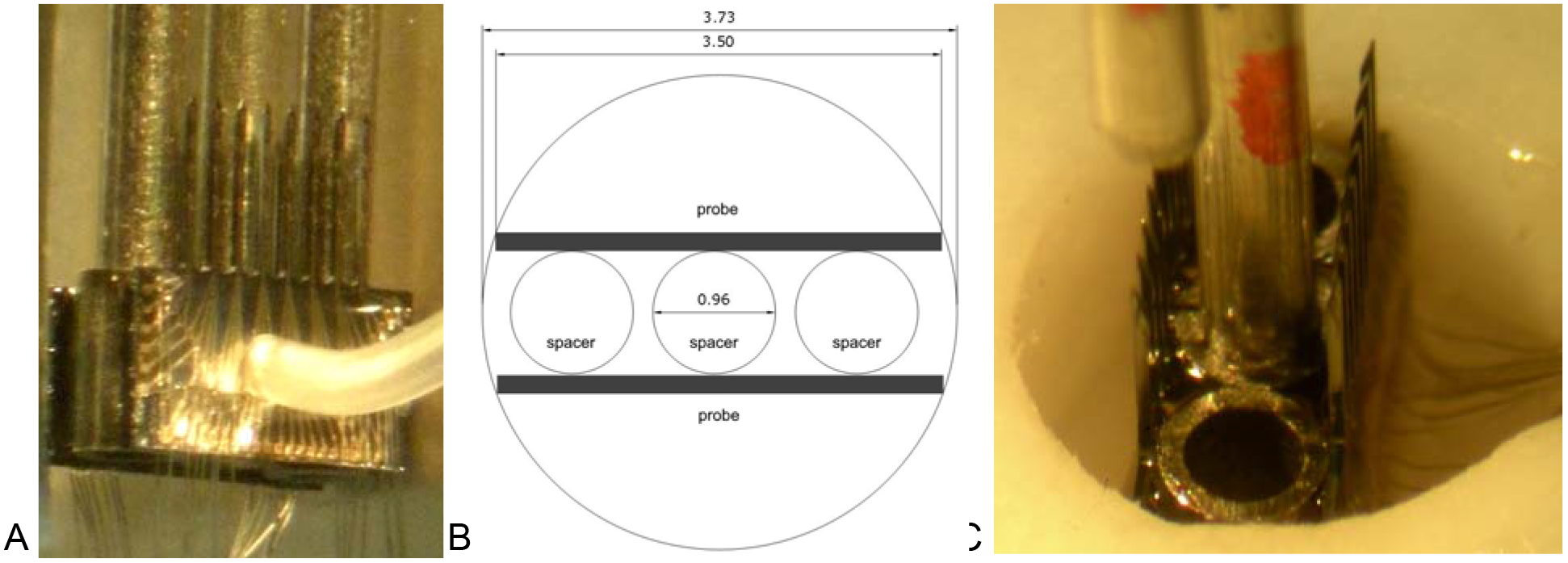
A) Photo of the array assembly with two planar probes and a 3-rod holder for suspending 3 spacers between the probes; B) top-view drawing of the molding chamber with the array and spacers (dimensions in mm); C) photo of the molding chamber with the array mounted on a micromanipulator.

### The array assembly

The array’s superstructure was protected from the saline environment by Parylene-C (as described above), silicone elastomer, and epoxy. Similar triple-layer polymer coating was used for protecting our earlier microwire and silicon-based arrays and demonstrated excellent long-term viability in vivo (Pikov, Bullara, and McCreery 2007). Before beginning the array assembly, the superstructure of each probe, including the demultiplexer and the bonding wires, was coated with a conformal layer of silicone elastomer (MED-4211, NuSil Technology, Carpinteria, CA) viscosity reduced with xylene to create an ion-impermeable protective conformal layer for active electronics (Song et al. 2005, Edell 2002). The assembly process was developed, as described below. For precise alignment of the probes, we used three spacers made from 18½ G stainless steel tubing (ID = 0.96 mm) with a height of 1.8 mm to match the height of the probe superstructure. A holder containing three rods made from 20 G stainless-steel tubing (OD = 0.81 mm) was inserted inside the spacers using a micromanipulator, glued to them with polyvinyl alcohol (PVA), and then lifted to keep the spacers suspended in the air, while two probes were pressed against them with Teflon tubing and bonded using a medical implant-grade (USP class VI) epoxy (EpoTek 301, Epoxy Technology) (Figure 3A). The epoxy was cured overnight in a 60°C oven. The holder with the attached array was then lowered into a round molding chamber (Figure 3B) to the depth of 2 mm below the surface, and two rods were retracted after dissolving the PVA with water, while leaving a single rod in the middle spacer (Figure 3C). The molding chamber was then filled with epoxy to about 1/3 of total height and cured overnight in a 60°C oven. The last rod was retracted after dissolving PVA with water, and the molding chamber was then filled with epoxy (EpoTek 310, Epoxy Technology) to the top and cured overnight in a 60°C oven. The array with the attached helical cable was then released from the molding chamber.

### Electrical testing

Prior to the implantation and at monthly intervals after the implantation, the electrical testing of the array was done to confirm the array and cable functionality. The testing was done by delivering a biphasic 10 μA current pulses and measuring the access impedance of each electrode site. The access resistance is the early rise in the voltage transient induced by the stimulus current pulses, it provides information about the electrode-tissue interface and the condition of the electrode site (Tykocinski, Cohen, and Cowan 2005). Any access impedance values > 200 kΩ indicated the electrode failure. For automating the access impedance testing, a 32-channel Reed relay switchbox (Model SeaI/O-440N, SeaLevel, Liberty, SC) was used for switching among 32 electrode sites; it was controlled by a custom Visual Basic program via the USB connection.

### Surgical procedures for array implantation

Young adult male cats weighing 2.5 to 4 kg were used. The animal studies were conducted according to the NIH guidelines and were approved by the HMRI Animal Care and Use Committee. Prior to surgery, the array and some surgical and testing tools (insertion tool with hex ranches, cable for perigenital stimulation, ball electrode for the recording of evoked potentials, and a connector cable for pressure transducer and a bladder catheter for urodynamics) were sterilized with ethylene oxide for 24 hours using a room-temperature system (Anprolene AN74i, Andersen Products, Haw River, NC). Other surgical tools were sterilized by autoclaving. The animals were anesthetized with isoflurane and nitrous oxide. Prior to the surgery, the cats were castrated to help prevent urethral blockage and to facilitate cleaning of the perigenital region. Two needles were inserted into the perigenital skin, 2 cm apart. The bladder catheter was inserted. The pressure transducer was filled with sterile saline and connected to the bladder catheter. The L_4_-S_1_ dorsal vertebral processes were palpated and the marked on the skin. The skin and underlying muscle were cut, and the L_6_ and L_7_ dorsal spinal processes were exposed and removed. The L_6_-L_7_ dorsal laminectomy was performed. Localization of the rostrocaudal spinal level involved in the control of micturition (typically S_1_ or S_2_ spinal segment) was accomplished by intraoperative recording of maximal evoked cord dorsum potential (ECDP) at the spinal cord surface with a ball electrode during electrical stimulation of the perigenital area at 20 mA, 0.2 ms/phase, and 10 Hz using a pair of needles (McCreery et al. 2004). The evoked potentials were recorded while moving the ball electrode rostrocaudally on the dorsal surface of the sacral cord in 1 mm increments. The DGC location was estimated to be at the maximal ECDP, which was defined as the maximal 2^nd^ peak of the evoked response. The dura over the DGC location was cut longitudinally at midline, the pial sheath was dissected over the dorsal roots, and the roots were gently retracted laterally to expose the spinal cord surface. The ECDP localization procedure was then repeated, while avoiding a confounding effect of stimulating the dorsal roots. The array was placed at a bottom of the barrel of our custom insertion tool (McCreery, Bullara, and Waldron 2001), with the vacuum applied through the barrel to keep the array in place. The barrel was positioned at the estimated DGC location using a stereotaxic manipulator with the side wings barely touching the cord to keep the dorsal roots retracted (Figure 4). The sensing probe located alongside the barrel of the inserting tool was used to detect an initial contact with the dorsal surface of the spinal cord, when lowering the barrel with loaded array. The electrode cable was routed rostrally and attached to the L_5_ process with a suture to stabilize the cable during the array insertion. Then, a stabilizing pad, made from polyester mesh and attached to the cable 10 mm from the array superstructure, was tamped onto the dura to limit the cable movement and reduce the transmission of torque and longitudinal pulling forces from the cable to the array (Figure 4).

**Figure 4.**
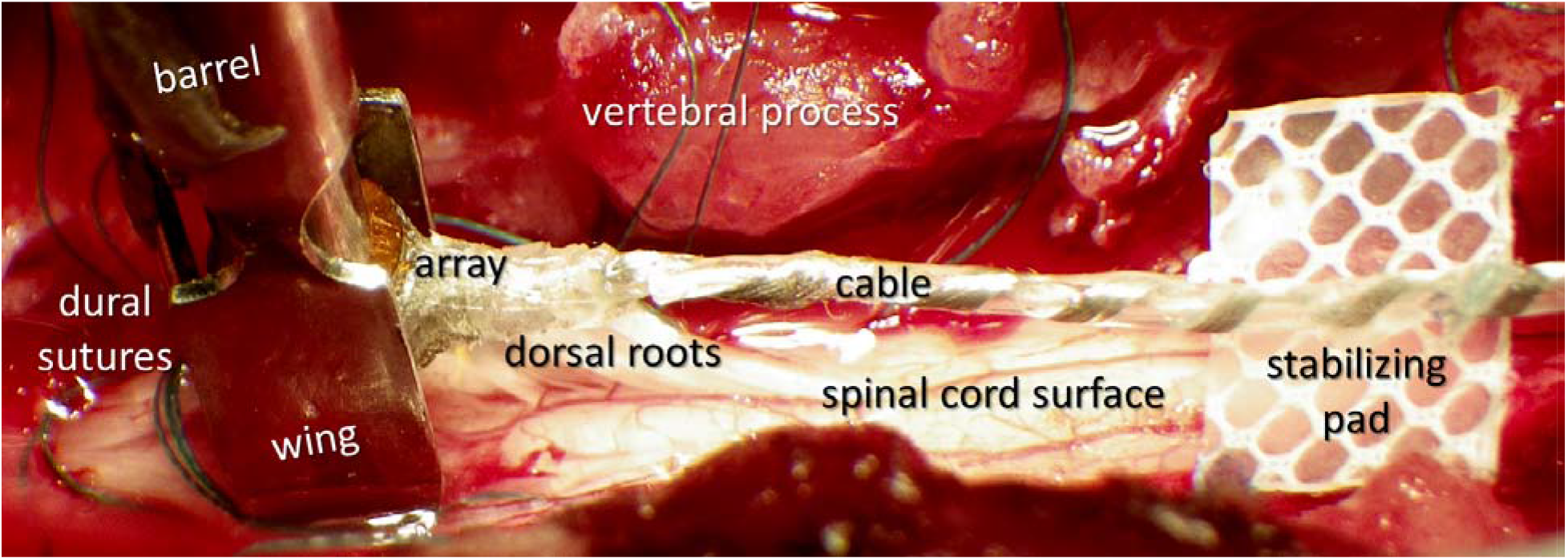
Photo of the spinal cord laminectomy with the barrel and wings of the inserter tool with a loaded array on the left and the stabilizing pad of the array cable on the right.

The inserter tool was deployed to inject the array into the spinal cord at a velocity of approximately 1 m/s and, due to its use of the sensing probe, its motion stopped just before the array superstructure impacted the cord surface. Completeness of the array insertion was confirmed using the surgical microscope with built-in video camera (S21, Zeiss) by observing the array superstructure resting directly on the pia. Next, absence of mechanical damage and completeness of insertion was confirmed by measuring the access impedance measurements from the shallowest (most dorsal) electrode site in each shank, as described in previous section. Large access impedance values (> 200 kΩ) indicated either the shank fracture or incomplete insertion, which was further distinguished by measuring the access impedance of the deeper-located electrode sites. After confirming the absence of mechanical damage and completeness of the array insertion, the dura was fully closed over the array with several 7-0 sutures. The electrode cable was tunneled subcutaneously to the skull using a guiding metal tube. The percutaneous connector was mounted on the skull with 4 stainless steel screws and bone cement (Methacrylate). The muscle, fascia, and skin were closed over the spinal laminectomy and around the percutaneous connector on the head. The animal was removed from the mounting apparatus and the surgical anesthesia wad discontinued. Intraoperative urodynamic testing was then performed while the animal was under light propofol anesthesia, as described in the next section. After the urodynamic testing, the bladder was emptied by draining the catheter to the collection bag. Once breathing on its own, the cat was placed in the heated incubator for recovery from anesthesia and monitored until regaining sternal recumbency.

During the first 3 postoperative days, cats received 0.01 mg/kg of analgesic Buprenorphine (s.c., every 6 h) or a transdermal Fentanyl patch (25 μg/hr) placed on a shaved area of the over the dorsal cervical region. Cats were attended daily for general inspection and cleaning the head connectors. Baytril (enrofloxacin, 22.7 mg tablets given orally) or Clavamox (amoxicillin, 62.5 mg tablets given orally) were given in alternation twice a day for 7-14 days after the surgery, as a prophylactic antibiotic. Appetite, behavior, urination, defecation, and any potentially pain-related behavioral signs, such as immobility or reluctance to move, abnormal posturing, decreased appetite, anxiety, aggression, and vocalization were monitored daily. For the first two weeks after surgery, animals were housed in a cage without perches to reduce spinal mobility and allow the array to become stabilized by the growth of connective tissue.

### Surgical procedures for spinal cord transection and animal care after the transection

The spinal cord transection (SCT) at T_12_ vertebral level was performed in one cat with good urodynamic responses to microstimulation at 45 days after the intraspinal array implantation. Three days prior to the SCT, blood work (complete blood count and clinical chemistry) was done to ensure that the blood parameters, and the kidney function specifically, were within normal limits. 24 hours before the SCT, the cat was given a health examination to check basic physiologic parameters (body weight, temperature, pulse, respiratory rate) and overall wellness of the animal (lack of abnormalities in the skin, superficial muscles, and bones). At that time, a transdermal Fentanyl patch (25 μg/hr) was placed on a shaved area of the over the dorsal cervical region. Food was withheld 12 hours prior to the procedure, but water was offered *ad libitum*. On the morning of the SCT, 20,000 U/kg of Penicillin G Procaine or 4-7 mg/kg of Enrofloxacin (Baytril) were administered i.m. and repeated 6-8 hours post-SCT. Ketamine, acepromazine, and atropine were administered i.m. as a preoperative anesthetic followed by administration of i.v. Pentobarbital. Maintenance anesthesia was isoflurane and nitrous oxide administered by inhalation. The spinal cord was exposed by a dorsal laminectomy. A dorsal midline skin incision was made from T_11_ to T_13_, the muscles were retracted, the T_12_ dorsal vertebral process was removed, and the laminectomy was performed. The dura was exposed for a length of about 15 mm. About 0.5 ml of 1% Lidocaine was topically applied and injected into the cord. The thin and blunt-tipped 5-mm-wide Teflon strip was passed under the cord. The ends of the Teflon strip were lifted slightly and the spinal cord was cut with dura-cutting scissors. The dura was then closed with 7-0 sutures and covered with a piece of Gelfoam to promote growth of connective tissue. The overlying fascia and skin were closed in layers. Post-SCT, meticulous care was given daily (7 days/week), including cleaning, urine expression (twice daily), exercising and massaging of paralyzed limbs to avoid joint fixation.

The cat was housed in a fiberglass enclosure with the floor measuring 3’ × 3’ lined with incontinence pads that were replaced several times a day as needed to keep them dry and prevent pressure sores and avoid soiling of the animal. To promote physical activity, the cat was exercised in specially constructed runs made from polyvinyl chloride tubing and polyethylene mesh and lined with foam matting enclosed in vinyl and covered with incontinence pads. Once a week, the run walls were dismantled and cleaned in the cage-washer or by hand steaming. Meticulous everyday care was provided, including cleaning, expression of urine and feces, and exercising and massaging of paralyzed limbs to avoid joint fixation. After the SCT, the cat did not clean or groom the parts of its body below the lesion, so the hindquarters were washed, dried, and brushed to substitute for normal grooming. Bladder catheterization or tactile induction of spontaneous bladder emptying was performed at least twice per day, and the urine was collected and tested using urine reagent strips (MultiStix 10 SG, Siemens Medical Solutions, Malvern, PA) for presence of blood and for possible urinary tract infection.

### Urodynamic assessment during microstimulation

In spinally-intact animals, the assessment of urodynamic responses to the intraspinal microstimulation was conducted at one-month intervals, while after SCT, the assessment was done at weekly intervals. The cats were anesthetized with Propofol, and the urinary bladder was catheterized. The low level of Propofol anesthesia was maintained with the animal breathing on its own and responding to strong sensory stimulation. This allowed monitoring of possible undesired effects of stimulation, such as hindlimb flexion, movement of the tail, or painful sensation. Hydrostatic pressure within the bladder vesicle was measured with a pressure transducer (Model 041500503A, Maxxim Medical, Athens, TX). The bladder was filled with sterile warm (37°C) saline to a bladder pressure of 15-20 mmHg, which is similar to a storage phase of the micturition cycle and is below the threshold to generate the reflex contractions of the urinary bladder. The constrictive force, or tone, within the EUS was measured with the second pressure transducer as an “infusion pressure”, the resistance to the infusion of saline through a port on the side of the catheter at a rate of 100 ml/hr. The EUS was localized by slowly moving the catheter along the urethra to the point of maximum infusion pressure. The data from the pressure transducer was amplified, digitized at 10 samples per second using a 12-bit data acquisition system (USB-6259, National Instruments), and displayed and stored on a computer using a custom Visual Basic program with the Measurement Studio ActiveX components. Selection of the electrode sites and the pulsing parameters was done using a custom Visual Basic program controlling digital and analog output channels of the data acquisition board (USB-6259, National Instruments). These output channels were used to drive the custom-built current-controlled stimulator. The cathodic-first biphasic pulses were applied to one electrode site at a time at a duration of 0.15 ms per phase, an amplitude of 50 μA, and a frequency of 20 Hz. The train of electrical stimuli was delivered for 30 sec, followed by a recovery interval of 60 sec before the next train. The charge density was maintained below 40 nC/phase and 2,000 μC/cm^2^ in order to prevent stimulation-induced neural injury in the feline sacral spinal cord (McCreery et al. 2004). Changes in the bladder pressure and EUS tone were calculated as a difference of average pressures measured during 30 sec of stimulation and during 30 sec immediately before the stimulation (the baseline period).

After measuring the EUS tone and bladder pressure response to microstimulation, the bladder was filled with saline warmed to 37°C to a pressure of about 30 mmHg, and the catheter was removed from the urethra for testing the effectiveness of the most effective electrode sites in inducing micturition. Voided fluid was collected into a plastic dish and its amount was measured with top-loading balances (Model PB-S, Mettler-Toledo, Columbus, OH). The voiding duration and voided amount were recorded. A new urethral catheter was re-inserted and the residual urine in the bladder was aspirated with a syringe and its amount was also recorded and used to evaluate the voiding efficacy. “Near-complete emptying” was defined the residual bladder volume ≤ 10 ml. The cats with spinal cord transection were also tested for voiding efficacy without anesthesia, since they do not have any sensations associated with the catheter insertion and bladder filling.

### Histology and image analysis of the spinal cord sections

At the end of the study, the animals were deeply anesthetized with Pentobarbital (Nembutal, 50 mg/kg, i.v.) heparinized (5000 units, i.v.), and sacrificed by transcardial perfusion through the aorta with a pre-wash solution (0.05% procaine hydrochloric acid in PBS) for 30 seconds, followed by the fixative (freshly prepared 4% paraformaldehyde in PBS) for 2 minutes. In five of the animals, the necropsy was performed to extract the spinal cord segments with implanted array for the histology, and the array position relative to the spinal cord level was validated by a complete dissection of the sacral spinal roots. The array was then carefully removed, and the spinal cord tissue was post-fixed overnight in 4% paraformaldehyde, dehydrated, and embedded in paraffin using the automatic tissue processing and embedding system (Autotechnicon Mono, SEAL Analytical, Mequon, WI). Transverse sections (perpendicular to the probe shanks) of the paraffin-embedded spinal cord tissue blocks were cut at a thickness of 6-7 μm using semi-automatic motorized microtome (HM 355, Thermo Fisher Scientific, Kalamazoo, MI). The sections were stained Cresyl Violet (Nissl stain) for defining the gray matter boundaries. Some sections were also immunostained with the antibody to neuron-specific nuclear protein NeuN (MAB377, 1:2K, Chemicon, Temecula, CA) and visualized using chromogen Vector Red (Vector Laboratories, Burlingame, CA). The stained sections were microphotographed using a 3-megapixel digital microscope camera (Spot RT, Diagnostic Instruments Inc, Sterling Heights, MI) mounted on the fluorescent microscope (BX41, Olympus, Cypress, CA). The transverse area of the spinal cord gray matter and spinal cord boundaries were averaged from multiple tracings of the spinal cord sections in these animals, as described in detail previously (Pikov, Bullara, and McCreery 2007). Using a custom image analysis program written in Visual Basic, we then determined the locations of 4 shank tips closest to the midline of the spinal cord as a function of distance from the central canal.

## Results

### The probe fabrication and array assembly

Based on our previous mapping study in the feline spinal cord (Pikov, Bullara, and McCreery 2007), the dorsomedial span of the DGC was determined to be from 0.7 to 1.6 mm from the dorsal surface (with the central canal located at 1.6 mm) while the lateral extent of the DGC was 1.0 mm from the cord’s midline. Therefore, the array of stimulating electrodes was designed to encompass an area of 0.9 mm dorsomedially and 2.1 mm bilaterally, with an equal 0.3 mm spacing of electrode sites in both dimensions. In the rostrocaudal dimension, two probes were spaced 1 mm apart, based on a precision of ~1 mm during localization of the rostrocaudal spinal level involved in the control of micturition by intraoperative recording of maximal ECDP in response to perigenital stimulation (McCreery et al. 2004). A minimal shank width was determined to be 50 μm to accommodate the electrode site width of 20 μm and three 3 μm-wide conductive traces spaced 2 μm apart. Due to quality issues experienced during microfabrication and probe processing after their release from the wafer, the probe yield was about 50% (based on visual inspection). For the array assembly from individual probes, we used only probes that passed the visual inspection. The assembly process was rather labor-intensive and complex, so over the course of the study, we made several procedural changes to reduce the labor effort and complexity, while improving the reliability. The Methods section above reflects the optimized assembly procedure.

### In vivo implantation

Eight cats were semi-chronically implanted with the intraspinal arrays for at least one month, with one cat implanted for 6 months. The probe thickness was increased to 50 μm compared to 15 μm thickness of the NeuroNexus probes implanted in our previous study (Pikov, Bullara, and McCreery 2007). There were no shank fractures during the high-speed insertion. In all animals, we tested the arrays without the demultiplexing ASIC chip, therefore each probe was connected using a 33-wire cable, with two cables running from the array subcutaneously to the cats’ head. In the first two animals, each cable had the wires encapsulated in silicone tubing with the OD of 0.2 mm, resulting in a rather bulky and inflexible cable with the OD of 10 mm. These cables caused repeated infections and skin erosions, associated with accumulation of the sero-sanquinous fluid along the cable. These infections and erosions were treated by flushing and draining the cable tract with saline and antibiotic solution, but ultimately, these animals had to be sacrificed after about one month of implantation. In subsequent six animals, we used cables consisting of 33 thinner wires with the OD of 0.063 mm, resulting in smaller cable OD of 4 mm. These cables and arrays remained electrically functional during the entire 4-6 months of implantation, as confirmed by periodic measurements of the access impedance of the electrode sites. As evident from Figure 5, there was a general trend toward gradual increase in the access impedance over time for the sites that remained below a failure threshold of 200 kΩ, likely due to fibrotic encapsulation of the probe shanks.

**Figure 5.**
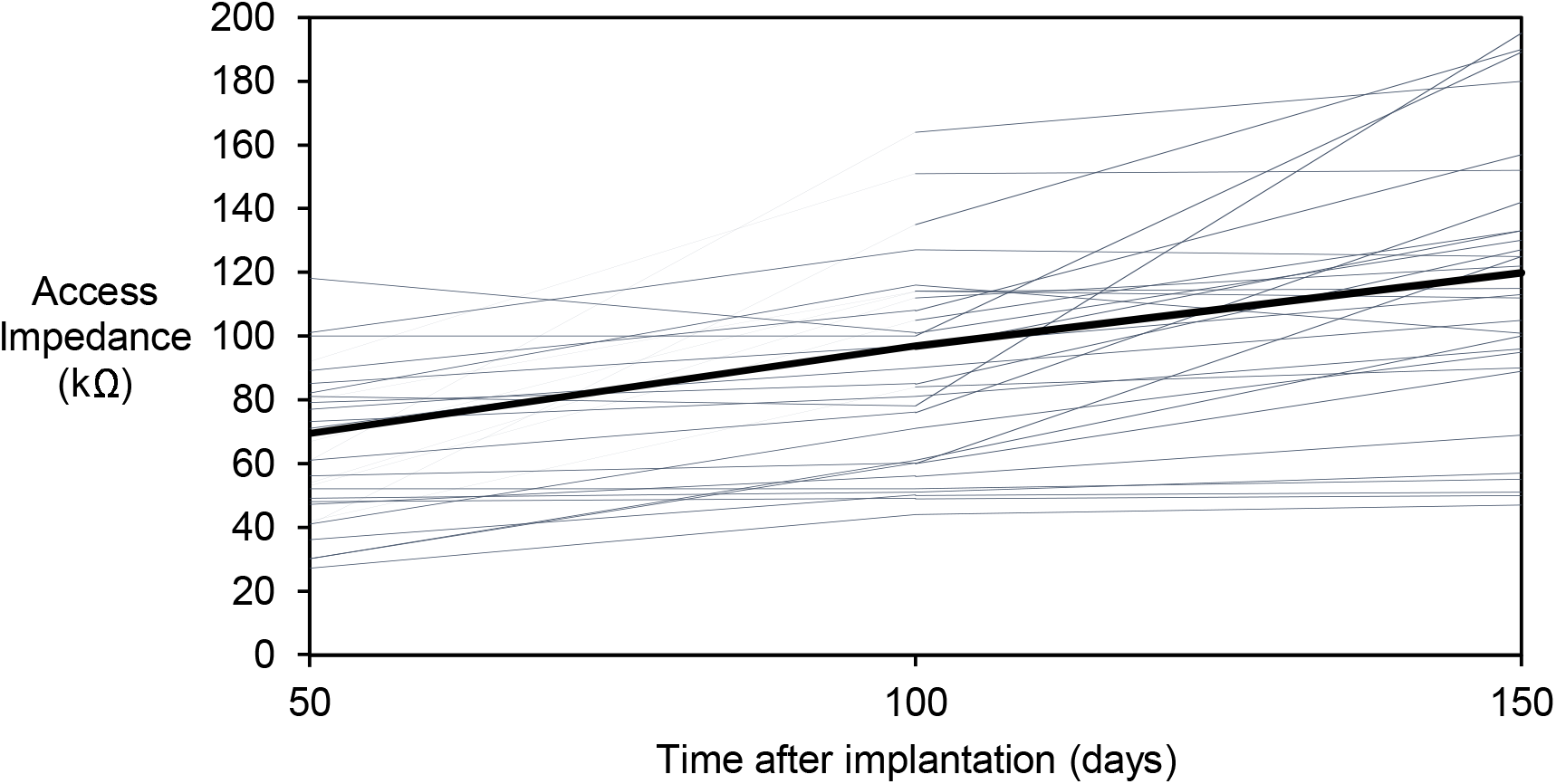
Access impedance of functional electrode sites (< 200 kΩ) over the course of implantation.

The effect of microstimulation on the bladder and EUS functions were evaluated urodynamically at about 1-month intervals. As shown in Figure 6 for one representative animal (SP07), the bladder responses to the electrical stimulation remained stable for a period of 5.5 months after implantation.

**Figure 6.**
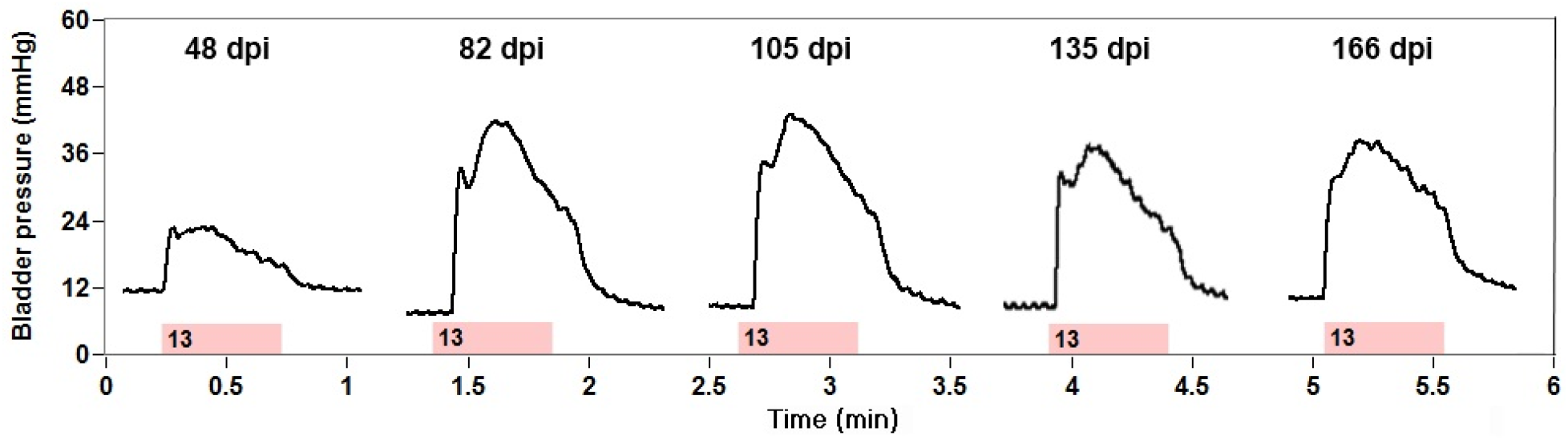
Stability of bladder responses to microstimulation over time after implantation. Abbreviations: dpi – day post-implantation; dpt – day post-transection. The pink bar indicates the duration of stimulation and the number inside the bar indicates the stimulated electrode.

In another animal (SP05), the bladder responses remained stable at 3.5 months post-implantation, therefore a low-thoracic SCT was performed at 4 months post-implantation to evaluate the effects of microstimulation in the absence of supraspinal control (Figure 7). There was an initial period of acute areflexia (days 1-5 post-SCT), requiring daily manual expressions of the bladder. By the day 7 post-SCT, the animal recovered spontaneous bladder contractions and was able to respond to microstimulation. By the day 22 post-SCT, the animal developed hyperreflexive non-voiding bladder contractions. At the days 44 and 58 post-SCT, the stimulation-induced contractions became stronger than pre-SCT.

**Figure 7.**
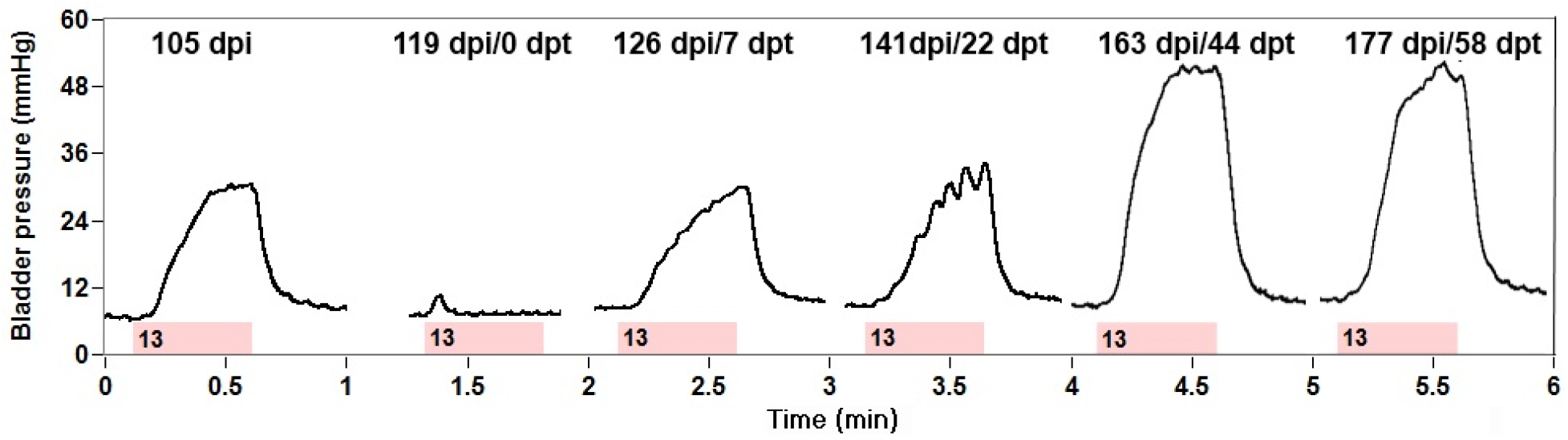
Bladder responses to microstimulation after low-thoracic SCT. Abbreviations: dpi – day post-implantation; dpt – day post-transection. The pink bar indicates the duration of stimulation and the number inside the bar indicates the stimulated electrode.

At the day 58 post-SCT in this animal (SP05), we also examined the effect of the stimulation on the bladder emptying by filling the bladder with saline and then removing the catheter to allow unobstructed flow of urine through the urethra. As seen on Figure 8, a stream of urine flow has been induced by stimulation of a single electrode site that was most effective in activation of the bladder and relaxation of the EUS. During 2.5 mins of electrical stimulation, the volume of voided urine was 77 ml, while the volume of residual urine was 27 ml, indicating a 74% voiding efficacy.

**Figure 8.**
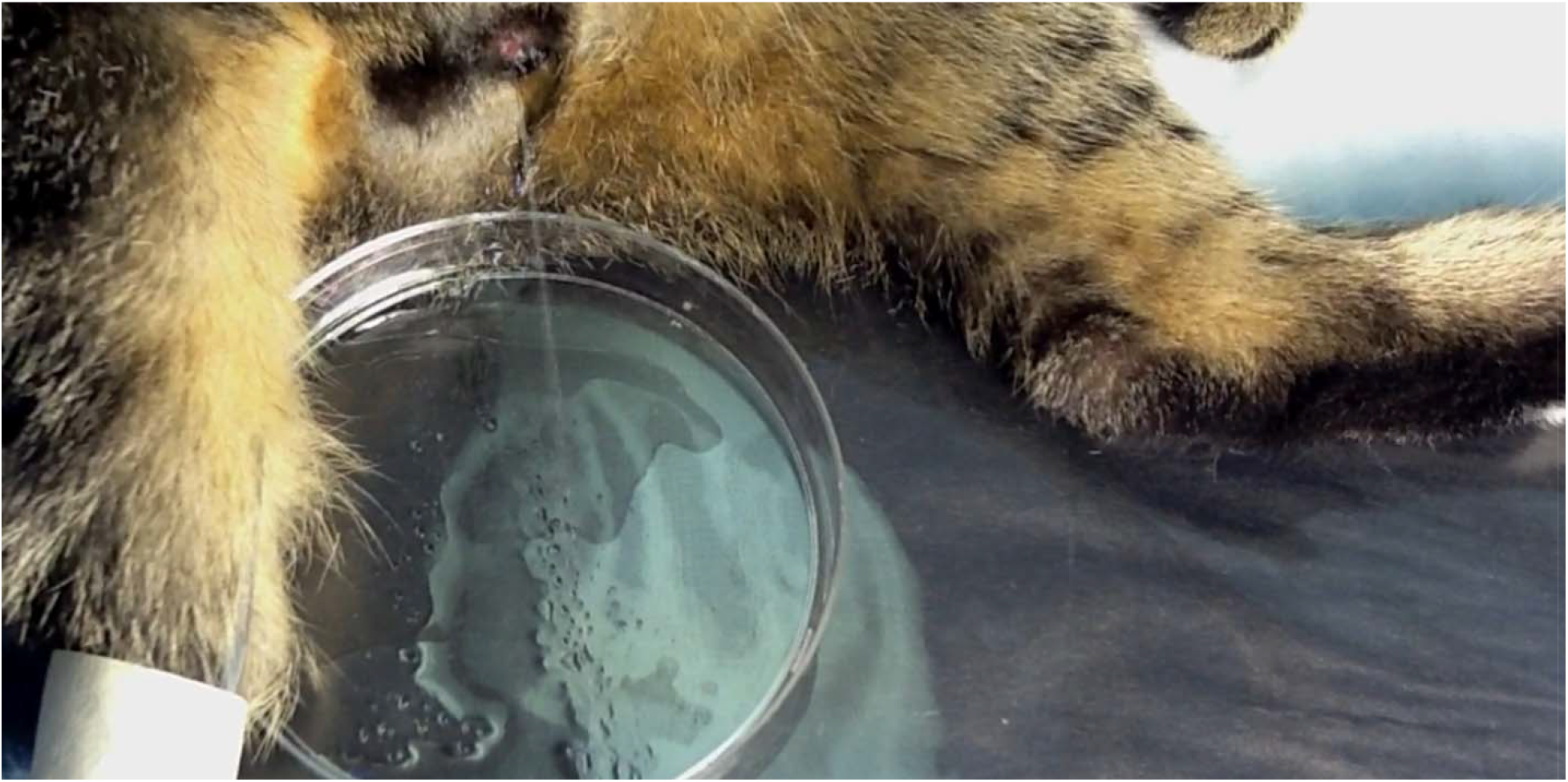
Bladder voiding induced by stimulating a single electrode site

After the animals were sacrificed, the necropsy and histology of the spinal cord was performed in five animals. Based on the dissection of the sacral spinal roots, we confirmed that the arrays in these animals were implanted at the rostral S_2_ segment of the spinal cord. As shown on the immunostained spinal cord section under the array (Figure 9), thickness of scar tissue around the DRIE silicon shafts was rather minimal (50-75 μm) and was similar to that of the scar tissue around NeuroNexus probes used in our previous study (Pikov, Bullara, and McCreery 2007), despite a larger cross-sectional area of the DRIE shanks.

**Figure 9.**
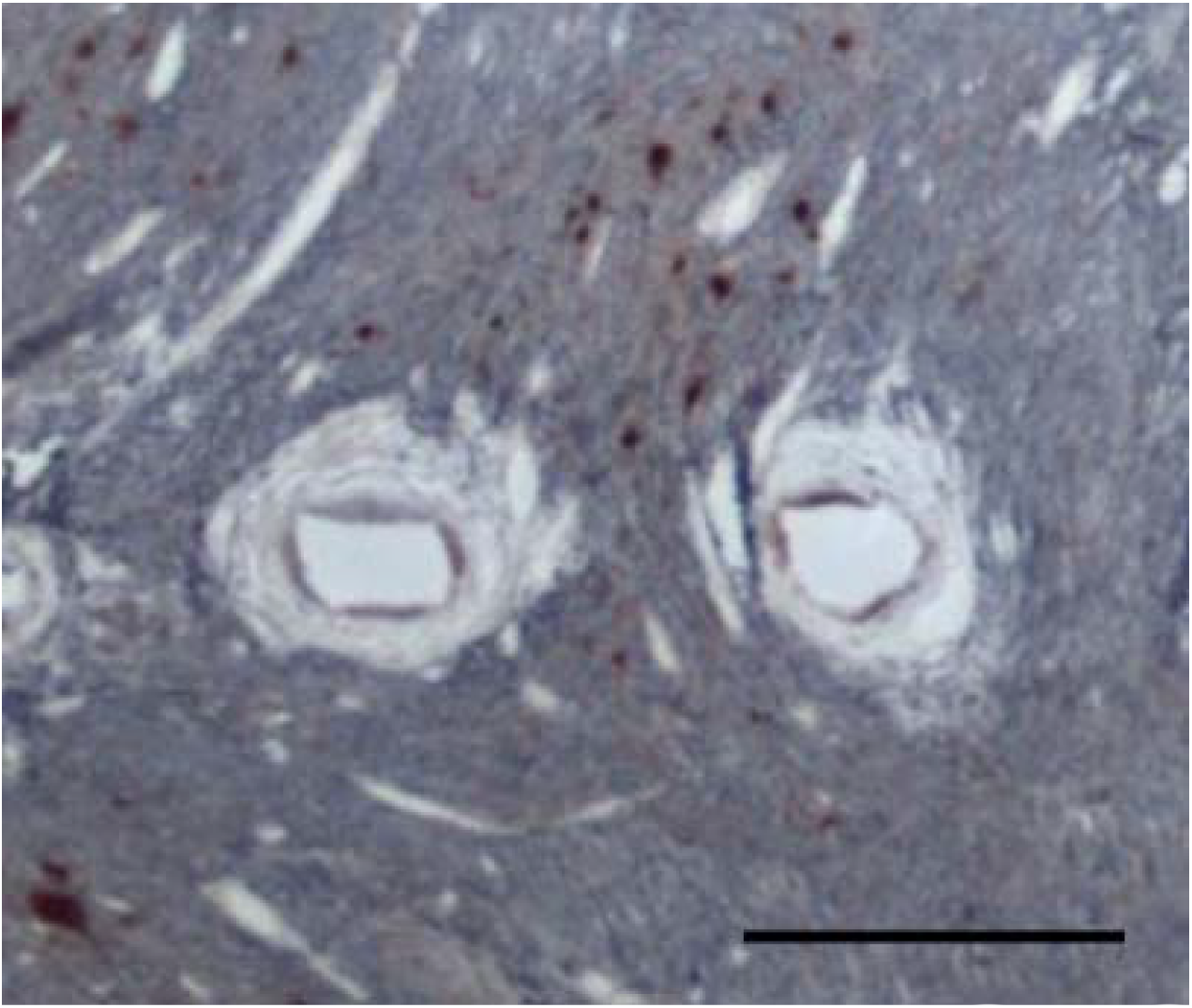
Photomicrograph of two probe shanks in the sacral spinal cord immunostained with NeuN and counterstained with Cresyl Violet. The scale bar is 300 μm.

The dorsal surface of the white and gray matter was considerably distorted by an indentation produced by a partial subsidence of the 1.8 mm-high probe superstructure, with a typical displacement of the spinal cord tissue being about 0.5 to 1 mm (data not shown). The image analysis was then performed to identify the locations of the stimulation sites on 4 shanks closest to the midline of the spinal cord. The near-midline shanks were selected due to: 1) their efficacy in eliciting micturition responses; 2) avoiding the variability in neuronal clustering near the lateral gray matter margin; and 3) for consistency in comparing with our previous mapping study that primarily utilized the single-site microwires (Pikov, Bullara, and McCreery 2007). In these 4 shanks from five animals, 26 sites (5.2 sites per animal), or 33% of the total number of sites induced elevation of bladder pressure (Figure 10). The effective and non-effective sites were co-localized, with no discernable sub-location within the DGC area exhibiting higher density of effective sites.

**Figure 10.**
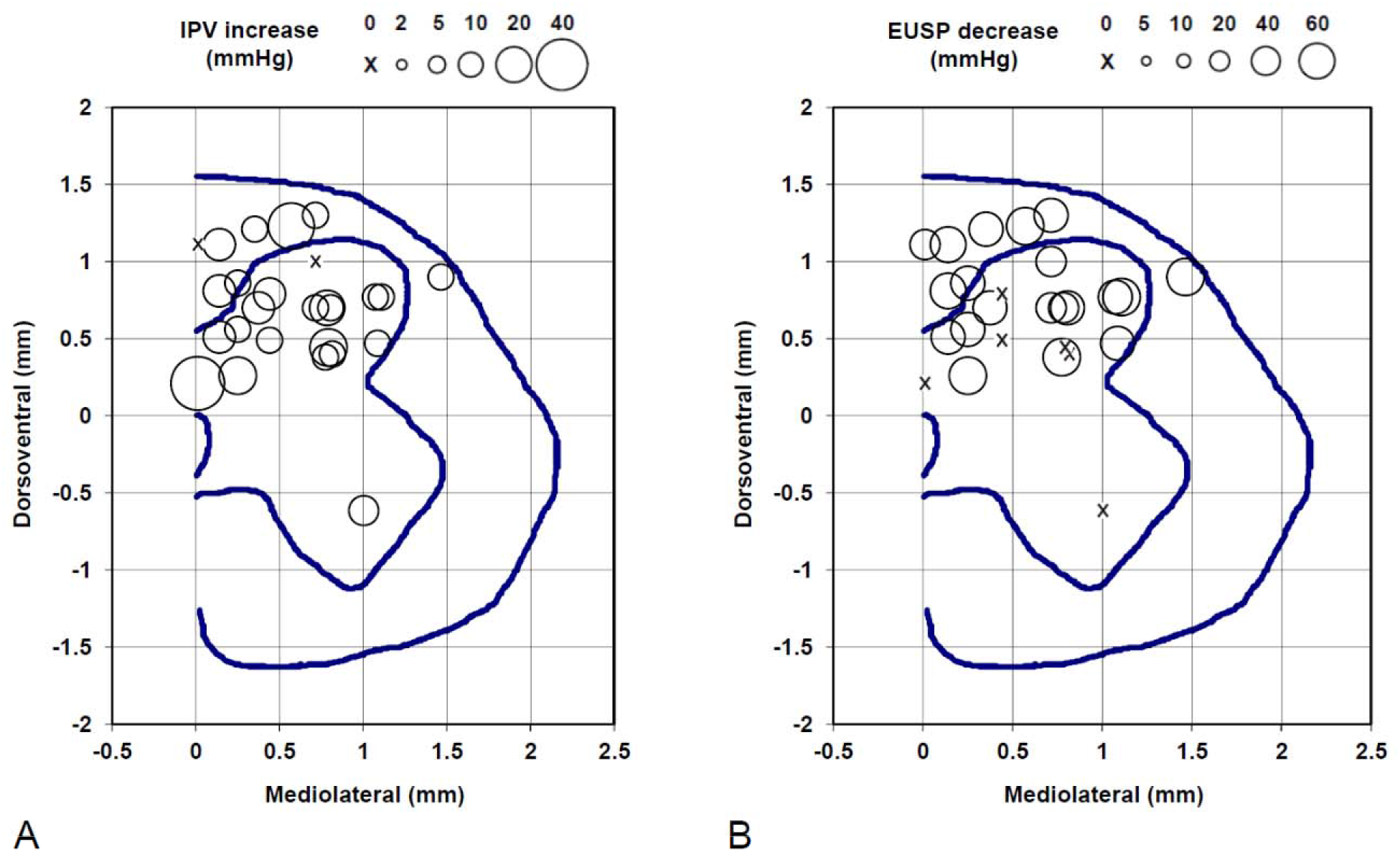
Locations of the electrode sites in the rostral S_2_ spinal segment that produced micturition-related responses: A) increases in the IVP and B) decreases in the EUSP. The circle size indicates the amount of IVP and EUSP change from the baseline level. The circle size scales are shown above the panels. The mediolateral and dorsomedial coordinates are provided in reference to the dorsal edge of the central canal. The blue lines in each panel represent the gray matter and the spinal cord boundaries and the location of the central canal.

## Discussion

This study confirmed and extended the findings of our previous spinal microstimulation study (Pikov, Bullara, and McCreery 2007). It was confirmed that the DRIE process was suitable for fabrication of chronically-reliable spinal stimulation probes and for a fracture-free insertion of the probes through the spinal pia in a moderately large animal (domestic cat), unlike the thinner silicon probes that require special mounting rigs during the insertion (Sakamoto et al. 2009). It was further demonstrated that the DGC area was involved in the coordination between the bladder and the EUS, as 33% of DGC-located electrode sites in the silicon-based arrays were effective in producing micturition-related responses, which is similar to 29% of effective sites observed with the single-site microwires (Pikov, Bullara, and McCreery 2007). In accordance with our previous study (Pikov, Bullara, and McCreery 2007), we also observed the increased effectiveness of stimulation in the DGC area for a period of 2 months after SCT, likely due to supraspinal disinhibition of the spinal central pattern generator circuitry for various functions (Pikov 2004, Ghali and Marchenko 2016) and/or hypertrophy and hyperexcitability of the spinal bladder afferents (Vizzard 2006, Yu et al. 2003, de Groat and Yoshimura 2009). In addition to taking the advantage of post-SCT amplification effects, the electrical stimulation in DGC can provide a more physiological approach of recruiting the bladder and EUS mononeurons, since the stimulation initially activates long-endurance/low-force type I motor units followed by incremental recruitment of short-endurance/high-force type IIb fibers (Hachmann et al. 2017).

The issues of glial scarring and foreign-body response to the implanted penetrating probes have been investigated mainly in the cerebral cortex. The spinal meninges are quite similar to the cranial meninges, but contain more collagen, which may serve as a mechanical reinforcer (Vandenabeele, Creemers, and Lambrichts 1996). Similar to the cortex, the electrode implantation causes a range of responses in the spinal meningeal layer, including fibrous tissue thickening (Eles et al. 2019), elastic softening (Moeendarbary et al. 2017), lymphocyte infiltration (McCreery et al. 2004), foreign-body response with microglial and astrocytic activation (Ersen et al. 2015, Kolarcik et al. 2012), and a risk of the cerebrospinal fluid leak (Short and Kirchner 1975, Miles et al. 1974, Nashold and Friedman 1972, Pineda 1978) and hematoma (Grillo, Henry, and Patterson 1974). In this study, the observed chronic foreign-body response was rather minimal, which was possibly due to a subdural placement of the array, stabilizing the array on the spinal surface with the stabilizing pad, and creating a thin uncoiled highly-flexible cable segment proximal to the array to promote its free untethered rostrocaudal movement together with the spinal cord inside the vertebral canal.

In this study, we observed considerable deformation of the spinal tissue following the array implantation. Similar spinal deformation was observed after epidural placement of electrode arrays made from shape-memory polymer and Parylene C (Garcia-Sandoval et al. 2018), polyimide and silicone (Minev et al. 2015, Capogrosso et al. 2018), and Parylene-C containing the electronics (Gad et al. 2013), as well as by penetrating microwire electrodes (Moritz et al. 2007, Bamford, Todd, and Mushahwar 2010). The compression force produced by the electrode array (especially, if tethered) may cause the fibrotic scarring and meningeal thickening, creating a physical barrier between the electrodes and the tissue resulting in increased activation threshold and vascular flow disruption. In extreme cases, the compression force can result in the array sinking into the tissue (Barrese et al. 2013, Maynard, Fernandez, and Normann 2000). In the future, the array design would be modified to reduce the height of its superstructure from 1.8 mm to <1 mm to reduce the downward force created by elastic dural encapsulation and a limited space inside the vertebral column.

Use of multisite probes with a large number of electrode sites created the challenge of interconnecting them to a bulky and relatively rigid cable with 33 wires, which resulted in repeated infections and skin erosions. That problem was addressed by designing the probe superstructure for interconnecting it with on-the-probe de-multiplexer ASIC chip which reduced the number of wires and by reducing the diameter of individual wires (along with their insulation). While the de-multiplexer approach was not tested during semi-chronic implantation, a reduction in the cable bulk and rigidity was successful in reducing the infections and skin erosions for the implantation period up to 6 months. For longer-term implantations, it would be beneficial to apply the de-multiplexer ASIC chip to further improve the cable flexibility.

Overall, over the course of iterative *in vivo* testing of several probe design options, we have arrived at an improved array design and confirmed its functionality during semi-chronic implantation. Further studies would be required to assess its longer-term reliability during one-year (or longer) implantations in preparation for eventual human translation of the intraspinal microstimulation array technology.

## Acknowledgments

This work was supported by the NIH grants R01-NS057287 (to VP) and R01-DC014044 (to MH) and the Department of Defense grant W81XWH-17-1-0538 (to MH). The silicon probes were fabricated by Kamal Yadev, tip-formed, cleaned, and wire-bonded by Vicki Cheng, and lased, assembled, and electroplated by Yelena Smirnova. Edna Smith assisted with the surgical procedures. Franchesca Sanchez and Jo Lemke performed the animal care. Nijole Kuleviciute performed the urodynamic testing and image analysis. Jesus Chavez performed the animal perfusion, necropsy, and histological work. The demultiplexer ASIC chip was provided by Prof. Wentai Liu at the University of California Los Angeles.

